# Assisted reproduction causes intrauterus growth restriction by disrupting placental lipid metabolism

**DOI:** 10.1101/030965

**Authors:** Yubao Wei, Shuqiang Chen, Xiuying Huang, Sin Man Lam, Guanghou Shui, Fangzhen Sun

## Abstract

IVF related intrauterus growth restriction or low birth weight (LBW) is very common in ART clinic. This study is focus on the aberrant lipid metabolism induced by in vitro fertilization and its mechanism. Firstly, we investigated the effect of IVF on fetal weight and placenta efficiency at E18.5 (at birth) and E14.5 (middle gestation). Data shows that IVF caused LBW and low placenta efficiency. Then we studied the lipidomics of E18.5 placenta and E14.5 placenta. The IVF group has an eccentric lipid content compared to *in vivo* group. All the 15 lipid classes are largely accumulated in E18.5 IVF placenta and are deficient in E14.5 IVF placenta. In detail, most of the 287 lipid species is accumulated at E18.5 and went short at E14.5. Using qRT-PCR we detected the expression level of genes related to lipid uptake, transport and metabolism. Most of these genes are down-regulated which indicated the metabolism function of placenta is disrupted seriously. To the imprinted genes for lipid metabolism regulation as *GNAS* and *Grb10*, IVF not only disrupt their imprinting status (methylation level) but also disrupt their gene expression. The expression of *DNMTs* and *Tets* are also disrupted in the placenta. These data demonstrate that IVF impaired the regulation network of lipid metabolism. These results prove the hypothesis: imperfect IVF condition of fertilization jeopardize the expression *DNMTs*, *Tets* and imprinting status of imprinted genes for lipid metabolism regulation. Then it causes to abnormal expression of genes for lipid metabolism and regulation. This leads to the significant differences in lipid species quantification and lipid metabolism. So it contributed to low lipid transport efficiency, restricted fetal growth and LBW. This study provides a renewed knowledge of lipid metabolism in placenta and its relation to imprinted genes and gave some clinical aware for optimizing the ART practice.

**Funding:** This work was supported by grants from the National Basic Research Program of China (973 Program) and National Natural Science Foundation of China.

**Competing Interests:** The authors have declared that no competing interests exist.

**Abbreviations:**

ART: artificial reproductive technology
IVF: in vitro fertilization
LBW: low birth weight
GNAS: Guanine Nucleotide Binding Protein Alpha
Grb10: Growth factor receptor-bound protein 10

## Introduction

Artificial Reproductive Technology (ART) is becoming the main technology for un-pregnancy couples in the last two decades, the total child born by ART is more than 5 million today [1, 2]. Children of certain percentage born by ART are with epigenetic phenotypes such as preeclampsia, low birth weight (LBW < 2,500g) [3], Angelman syndrome, Beckwith-Wiedemann syndrome [4], etc. And they in adulthood, they have a more inclined to chronic diseases like diabetes, hypertension, coronary artery disease [5], compared to natural born individuals. The different between ART and natural born is in vitro fertilization (IVF) and embryo culture for 3-4 days. So IVF will consistently affect the embryo development. The aberrations could be measured by methylation status of imprinted genes [6] or other indicators. To the LBW in clinic, the singleton infants by ART is 2.6 fold of infants naturally conceived [7].

Placenta plays a vital role for fetal nutrition and growth at post-implantation stage. Lipid metabolism is the main function of placenta. The lipid metabolism is vital for fetal and placenta development [8]. If lipid metabolism is in dysfunction, the embryo will growth slowly even death and result in slow fetal growth or LBW. As the technology advancing, Lipidomics could give us comprehensive information of lipid species and quantities in placenta.

There are more than 30 imprinted genes are play a role in the regulation of lipid metabolism. Imprinted genes have conservative functions for placental mammal animals. There are about 210 imprinted genes for human and about 130 for mouse [9]. Almost half of them express specifically in placenta. Knockout mouse models show that imprinted gene loss function will result in growth retardation or embryo lethal.

To investigate the lipid metabolism and its association with imprinted genes placenta, firstly, we validated the low placenta efficiency (or LBW) in mouse at E18.5 (later gestation) and E14.5 (mid-gestation), then we carried out the lipidomic analysis of mouse placenta. Secondly, we test the expression of genes for lipid uptake, transport and metabolism. Thirdly, we test the expression of genes for lipid metabolism regulation at RNA level and protein level. Fourthly, we study the methylation status and gene expression level of imprinted genes which participated in lipid metabolism. Then we test the gene expression and protein level of DNMTs and Tets which maintain the imprinted status of imprinted genes. The results demonstrate the aberrant imprinting status is associated with mis-regulation of lipid metabolism, which in turn causes abnormal expression of genes for lipid metabolism, such as lipid accumulation and transportation. And thus it contributes to low lipid transport efficiency leading to restricted fetal growth and LBW. Our study provides new insight as to improve IVF practice. To our knowledge, these results provide the first evidence of lipidomics of placentae and its association with imprinted genes.

## Author Summary

Birth weight is an important marking linked to adulthood health and diseases. Epidemiological studies indicated that low birth weight (LBW) is a common phenomenon among children conceived by in vitro fertilization (IVF), but the mechanism of LBW-associated with IVF is still unclear. Our lab’s recent work has shown that IVF can cause mal-development and dysfunction of the placenta. This study tests our hypothesis that IVF may cause LBW by disrupting lipid metabolism in the placenta. By comparing the weight of placenta and fetus during gestation and at birth (E18.5) between IVF and in vivo group, we found that IVF resulted in reduction of both fetal weight and placenta efficiency. Lipidomic analysis shows that all the 15 lipid classes were largely accumulated in IVF placenta at birth. Using qRT-PCR and Mass spectrum, we show that IVF disrupted the expression of genes related to lipid metabolism. The expression level of lipid regulation genes such as *Mtor, HMOX1* were also altered by IVF. Moreover, we have identified that imprinted genes such as *GNAS*, *Grb10* are important for lipid metabolism regulation; both their expression and imprinting status (methylation level) in the placenta were disrupted by IVF. Our data support the idea that IVF could disrupt lipid metabolism in the placenta by altering imprinting status of imprinted genes, and expression level of genes involved in lipid metabolism, which in turn contributes to reduced intrauterine fetal growth.

## Results

### 1 IVF results in intrauterus growth restriction

Placenta efficiency is a convenient index to measure placenta function. The calculation is fetus weight divided by placenta weight. We collected mouse fetuses and placentae at E18.5 and at E14.5 respectively and then measured their weights (Figure 1). At E18.5, placentae from the IVC, IVF groups were found to be significantly heavier than those of the *in vivo* group. The fetal weight of IVC and IVF groups was significantly lower than *in vivo* group (p < 0.05). There is no significant difference between the IVC and IVF groups. The IVF placenta weight were significantly heavier than the IVC placentae (p < 0.05), suggested that IVF itself can contribute to abnormal placenta development. At E14.5, there were no differences between placental weight but fetal weight of both the IVF and IVC groups was significantly lower than *in vivo* group (p < 0.05) (Figure 1). So ART treatment can disturb mouse placental and fetal development at gestation stage.

**Figure 1.**
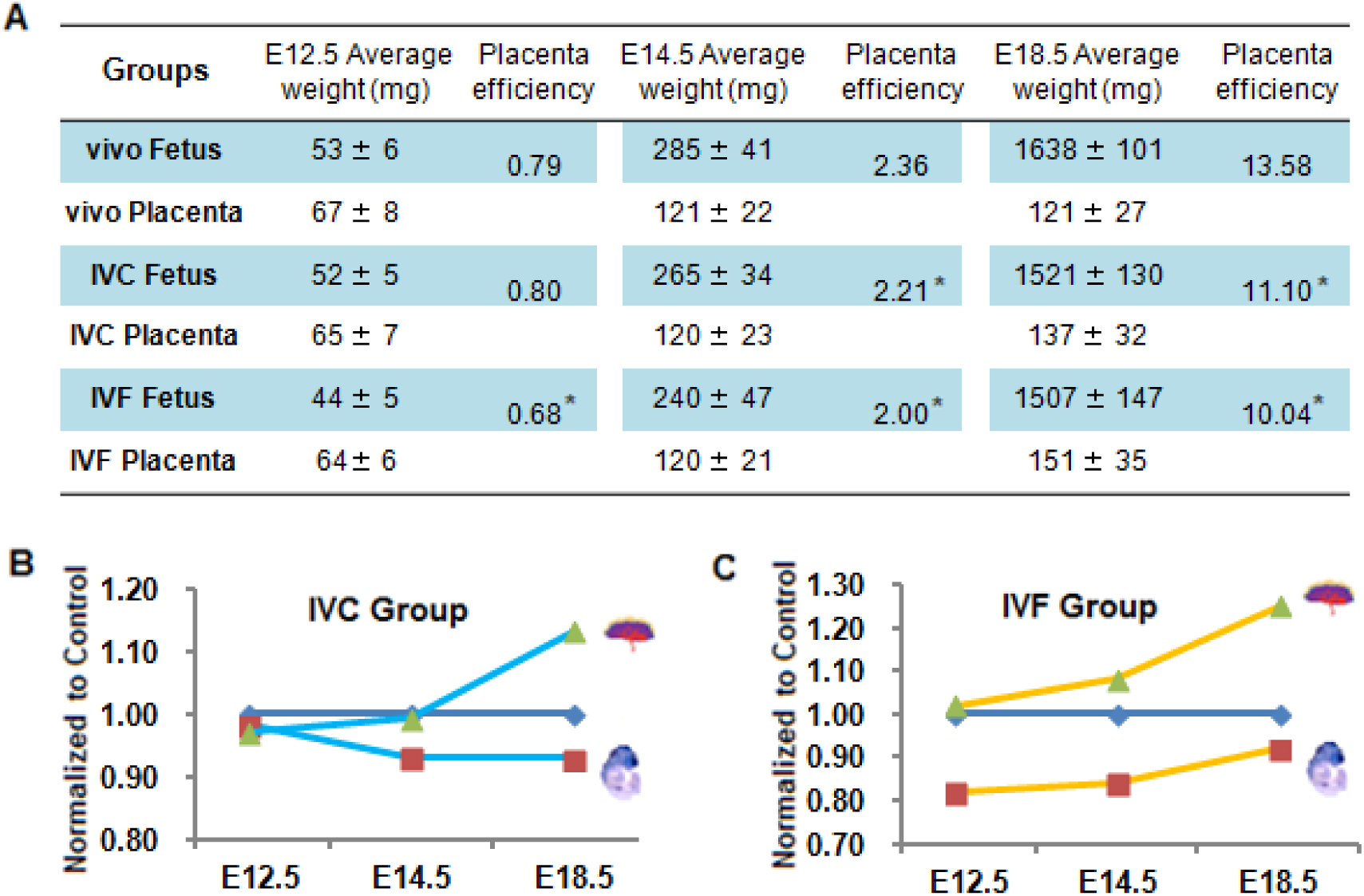
Fetal weight, placenta weight at E18.5 and its normalization of IVF results to the control. **A.** Average fetal weight, Average placenta weight and placenta efficiency are of *in vivo*, IVC, IVF groups. Data was showed as mean ± deviation. At E12.5, in vivo n=37, IVC n=40, IVF n=39. At E14.5, in vivo n=42, IVC n=38, IVF n=41. At E18.5, in vivo n=40, IVC n=36, IVF n=42. At E18.5, in vivo n=40, IVC n=36, IVF n=42. **B.** Normalization of IVC fetal weight, placenta weight to control (*in vivo*). The trend for fetal weight is lower to control, and for placenta weight is higher. **C.** Normalization of IVF fetal weight, placenta weight to control (*in vivo*). The trend for fetal weight is lower to control, and for placenta weight, it becomes more and more high.

### 2 Lipidomics show IVF placentae have abnormal lipid accumulation

Lipidomics is an efficient way to analysis the lipid species and content of animal tissues. We prepared E14.5 and E18.5 placenta sample and carried out the lipidomic analysis. There are 15 lipid classes were found in the placenta: Free Cho (free cholesterol), PC (phosphatidylcholine), SM (sphingomyelin), PE (phosphatidylethanolamine), PS (phosphatidylserine), PI (phosphatidylinositol), TAG (triacylglycerides), CE (cholesteryl esters), LPC (lysophosphatidylcholine), Cer (ceramide), DAG (diacylglycerol), GluCer (Glucosylceramide), PG (Phosphatidylglycerol), GM3 (anglioside mannoside 3), LBPA (Lysobisphosphatidic acid) (Figure 2A). The three lipid classes Free cho, PC, SM take about 2/3 of the total lipid mass of placenta (Figure 2B, C). To LPC, Cer, DAG, GluCer, PG, GM3 and LBPA, their content are less than 0.5 uM/g in the placenta. The result of the 15 classes in IVF group is significant (p < 0.05) higher than in vivo group at E18.5 (Figure 2). At E14.5, the content of 15 lipid classes in IVF group is less than in vivo group or IVC group (Figure 3).

**Figure 2.**
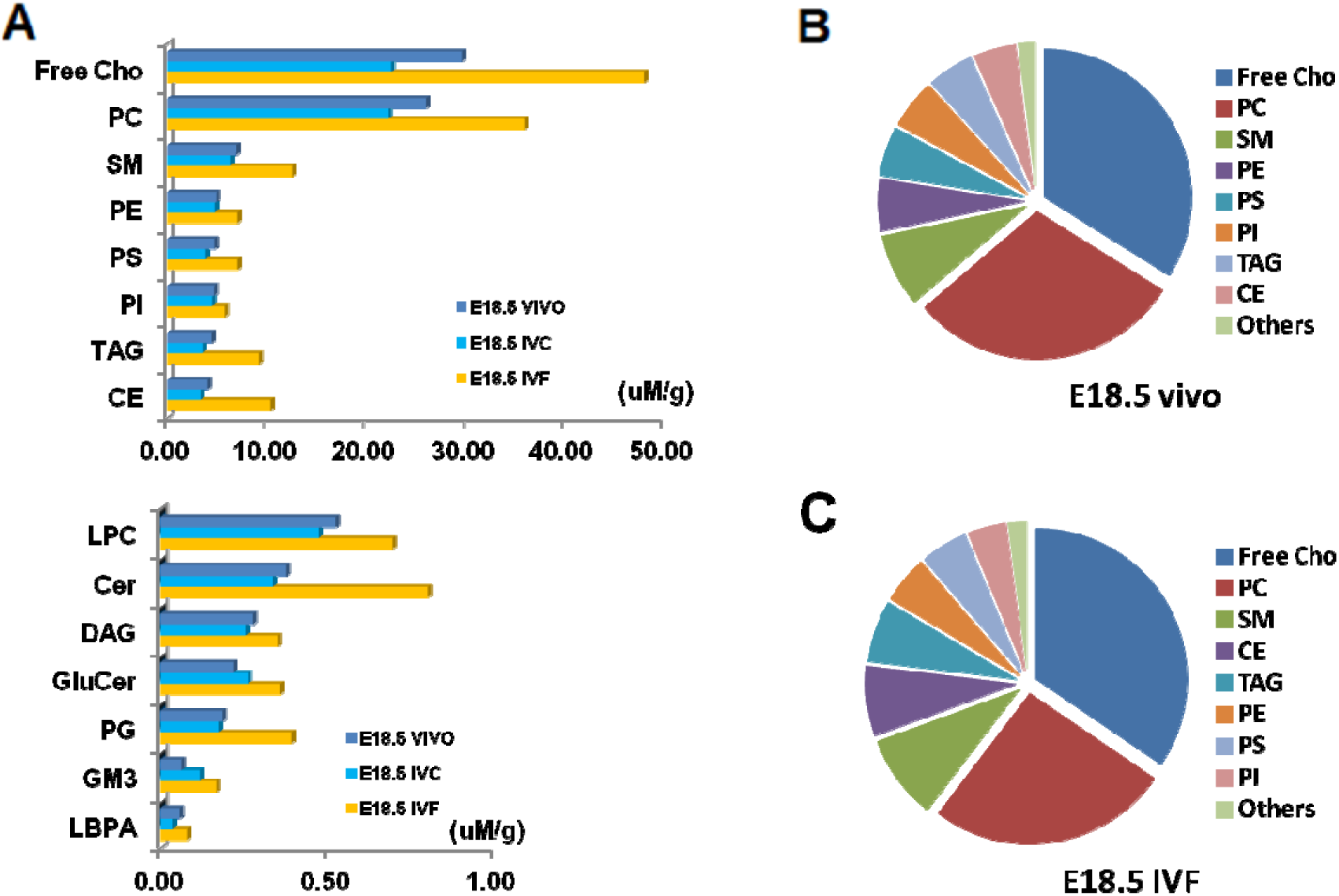
Comparison of 15 lipid classes in the placenta derived from in vivo, IVC or IVF at E18.5. **A.** The absolute content of 15 lipid classes as Free Cho, PC, SM, PE, PS, PI, TAG, CE, LPC, Cer, DAG, GluCer, PG, GM3, LBPA in the E18.5 placentae of the three groups. **B. C.** Relative content of the lipid classes in E E18.5 placentae of *in vivo* and IVF group (The proportion for IVC is about 60%).

**Figure 3.**
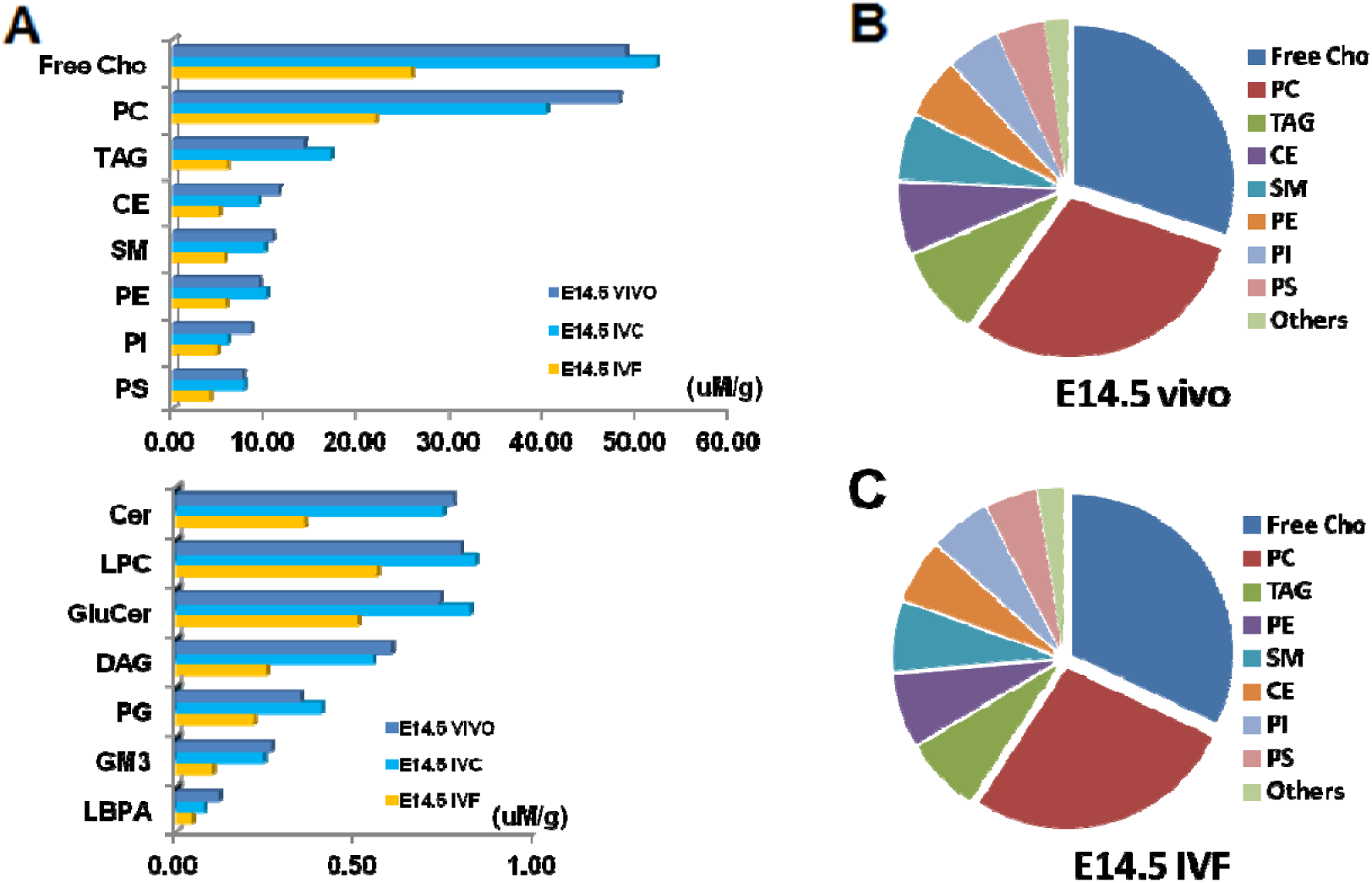
15 lipid classes in *in vivo,* IVC, IVF placentae at E14.5. **A.** The absolute content of 15 lipid classes as Free Cho, PC, SM, PE, PS, PI, TAG, CE, LPC, Cer, DAG, GluCer, PG, GM3, LBPA in the E14.5 placentae of the three groups. **B.** Relative content of the lipid classes in E E14.5 placentae of *in vivo* group; the mass of Free Cho and PC takes a proportion as 60%. **C.** Relative content of the lipid classes in E14.5 placentae of IVF group; the mass of Free Cho and PC takes a proportion as 59%.

The 15 lipid classes are composed by 287 lipid species. We make a heatmap by the relative content to in vivo group (Figure 4). For IVF group, almost every species is less than in vivo group at E18.5 and less than in vivo group at E14.5. The heatmap range is about 0.2 to 2.7 fold for different lipid species. The results indicate lipid mass uptake from mother is accumulated at E18.5 placenta but is in shortage status at E14.5 placenta.

**Figure 4.**
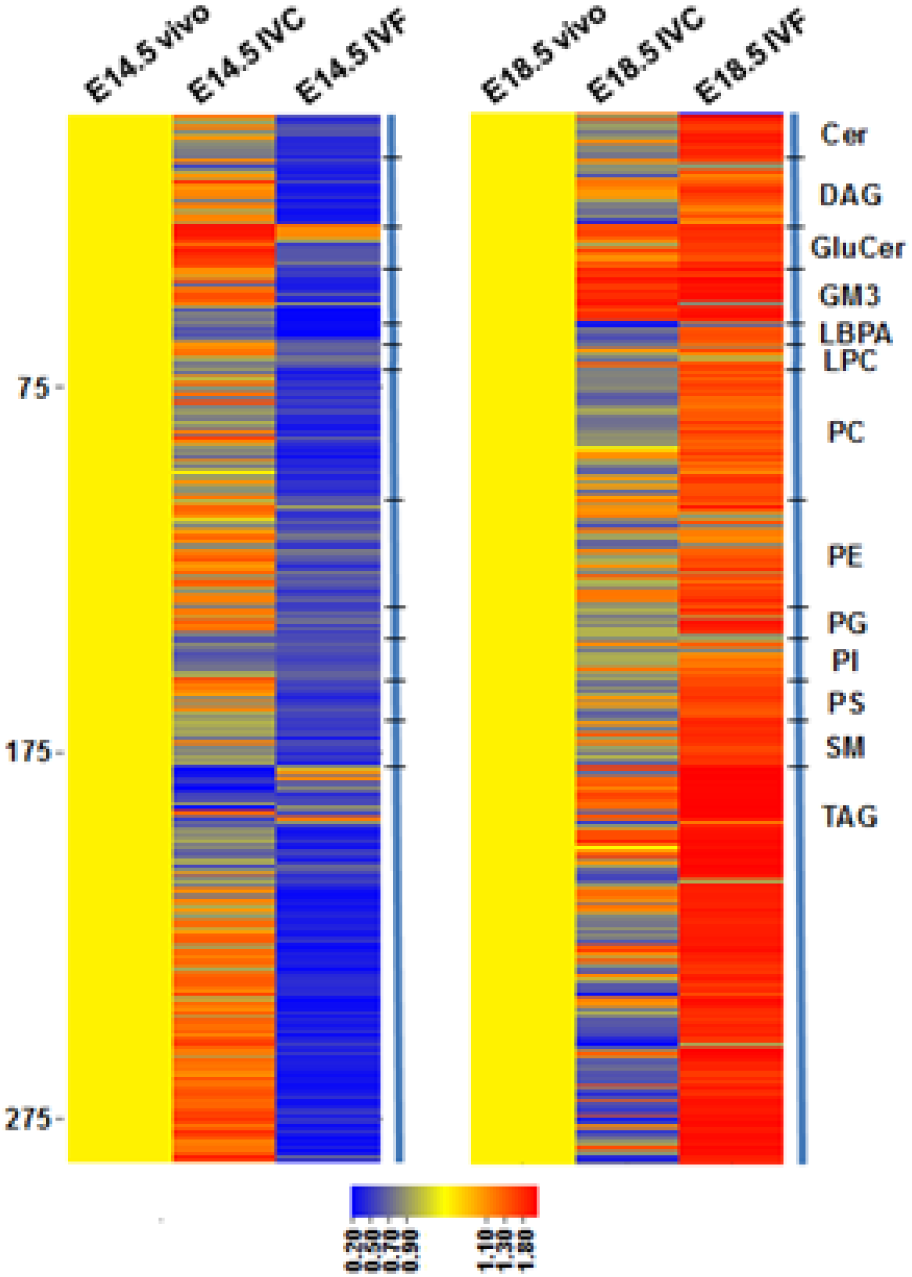
Heatmap of the 287 lipid species in placentae of the three groups

### 3 IVF disrupted the expression of genes for lipid metabolism in placentae

The lipid function of placenta are composed of lipid uptake, lipid transport and lipid metabolism. So how IVF affected the gene expression related to the three aspects is a key question. We used qRT-PCR to get the expression level of these genes. Compared to in vivo group, *CTPS1, PPARG, RBP1, SLC2A1* and *UGT1A2* of IVF group were significantly (p < 0.05) down-regulated at E18.5, *UGT1A1* was significantly (p < 0.05) up-regulated at E18.5 (Figure 5A). At E14.5, *UGT1A1, UGT1A2* were significantly (p < 0.05) down-regulated, *CTPS1, PPARG* and *UGT1A6* were significantly (p < 0.05) up-regulated (Figure 5B).

**Figure 5.**
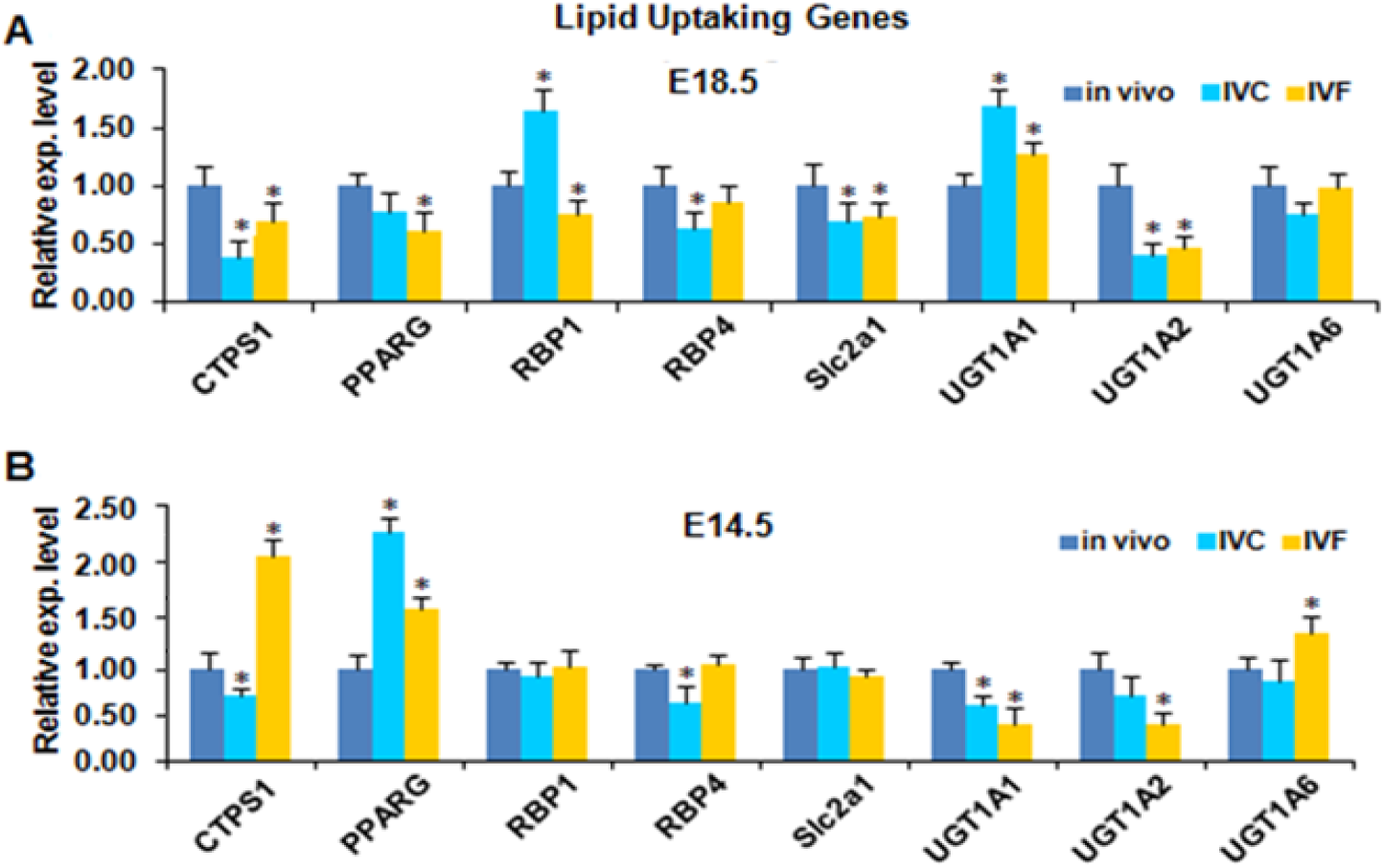
IVF altered the expression of lipid uptake genes in placenta. **A.** The gene expression levels of lipid uptake genes of the three groups at E18.5. **B.** The gene expression levels of lipid uptake genes at E14.5. Values expressed as means ± standard errors. * p < 0.05 compared with control.

For lipid transport genes, expression level of *LDLR, LRP1, LRP8, PLA2G15, FAT1* and *VLDLR* of IVF group were significantly (p < 0.05) down-regulated at E18.5, *APOE* and *LRP2* were significantly (p < 0.05) up-regulated at E18.5, compared to *in vivo* group (Figure 6A). At E14.5, *APOE, LDLR* and *LRP2* were significantly (p < 0.05) down-regulated, *LRP2, LRP8, PLA2G10, FAT1* and *VLDLR* were significantly (p < 0.05) up-regulated (Figure 6B).

**Figure 6.**
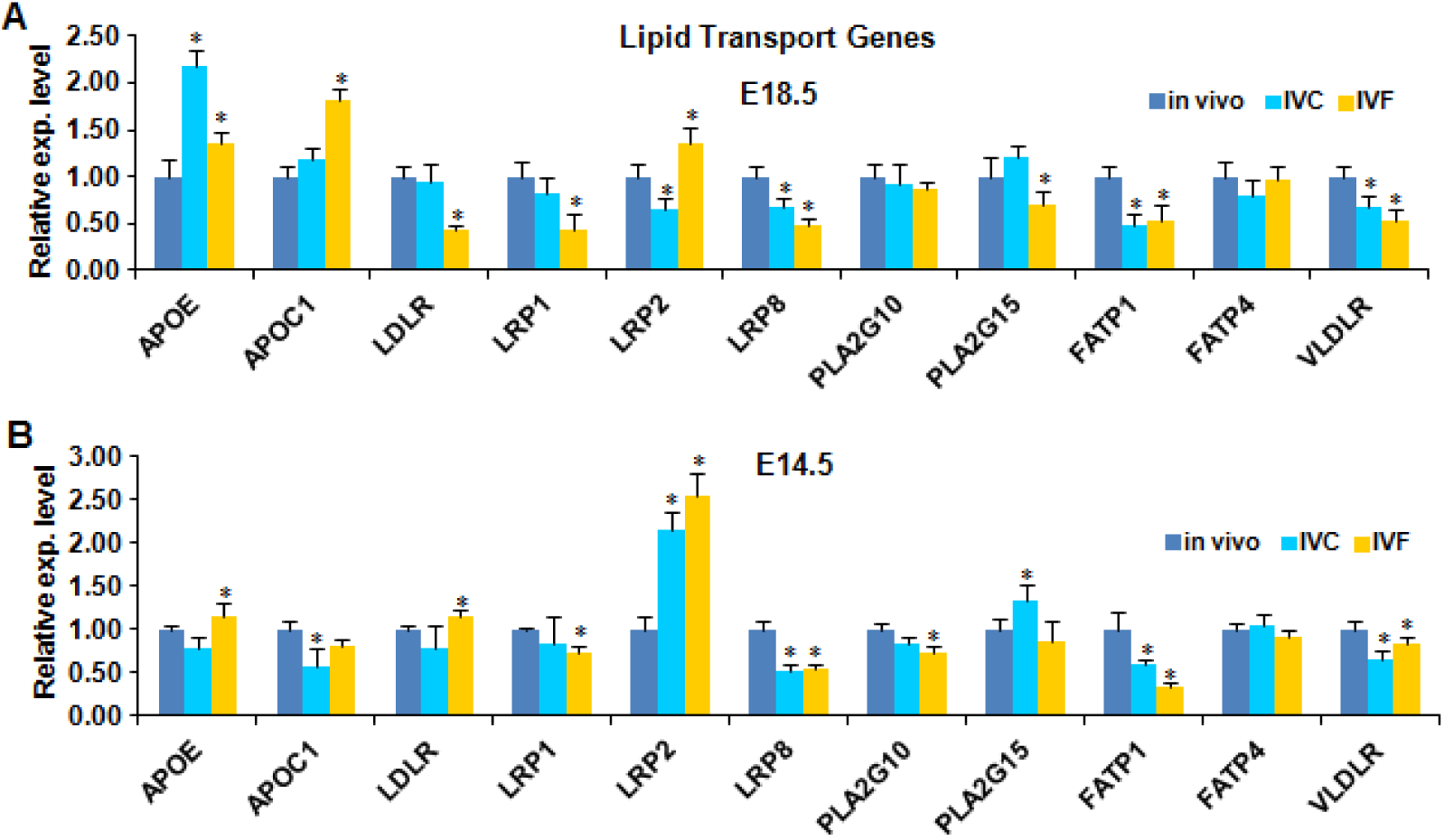
Effect of IVF on expression of lipid transport genes in placenta. **A.** Comparison of gene expression levels of lipid transport genes of the three groups at E18.5. **B.** Comparison of gene expression levels of lipid transport genes at E14.5. Values expressed as means ± standard errors. * p < 0.05 compared with the control (in vivo).

For lipid metabolism genes, expression level of *ABCG1, CTP1A, EL, FASN, LIPIN1, LIPIN3* and *LPL* of IVF group were significantly (p < 0.05) down-regulated at E18.5, *APOC1, CTP1B* and *PPARA* were significantly (p < 0.05) up-regulated at E18.5 (Figure 7A). At E14.5, *ABCG1, ACLY, EL, PLIN2* and *PPARD* were significantly (p < 0.05) down-regulated, *CTP1A, CTP1B, FASN, HSL, LIPIN1, PPARA* and *SEPP1* were significantly (p < 0.05) up-regulated (Figure 7B). So IVF seriously affected the expression of genes related to lipid uptake, transport and metabolism in mouse placenta.

**Figure 7.**
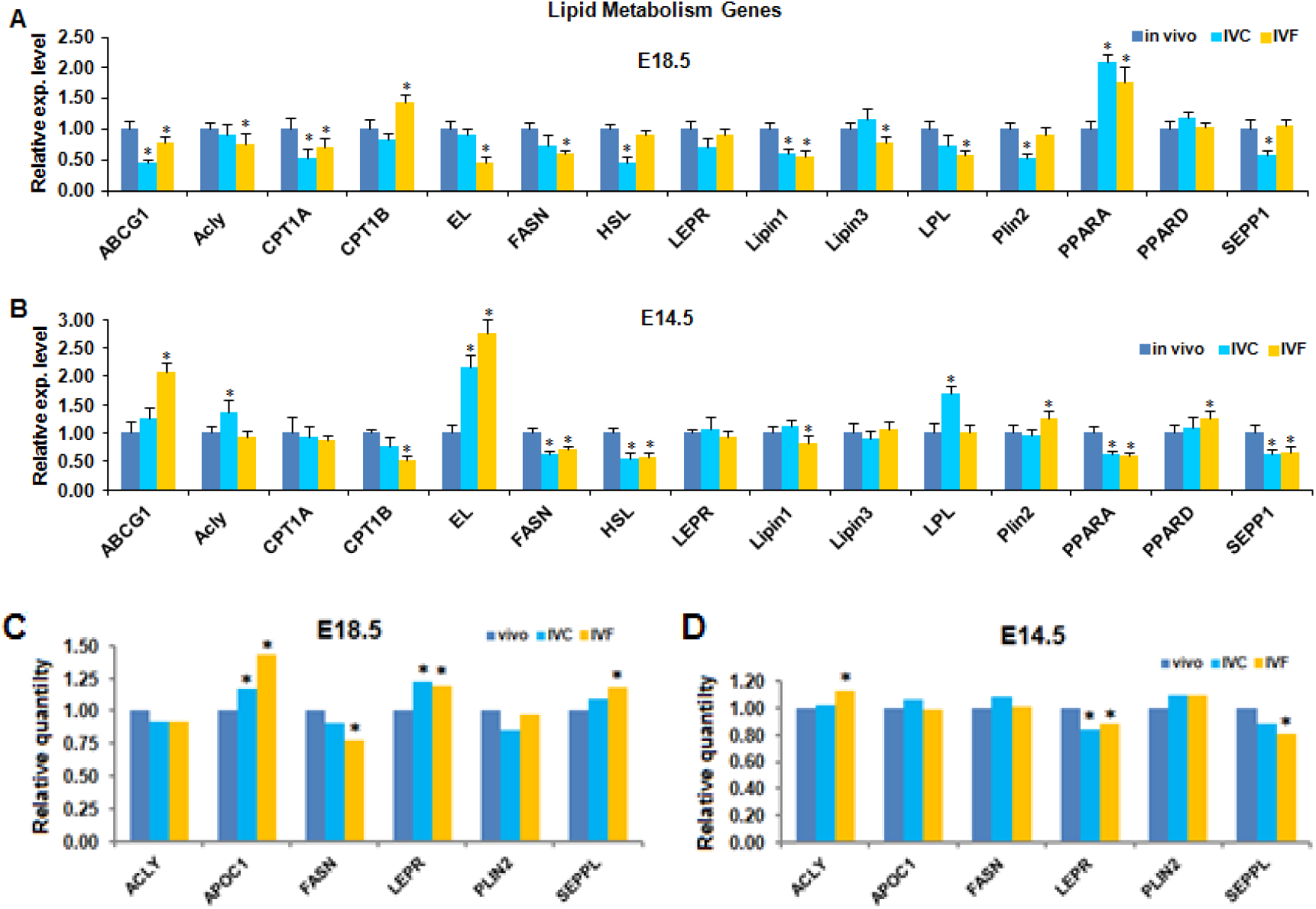
Effect of IVF on expression of lipid metabolism genes in placenta. **A.** The gene expression levels of lipid metabolism genes of the three groups at E18.5. **B.** The gene expression levels of lipid metabolism genes at E14.5. **C.** Comparison of the protein quantity at E18.5 placenta of the three groups. **D.** Comparison of the protein quantity at E14.5 placenta. Values expressed as means ± standard errors. * p < 0.05 compared with control.

### 4 IVF disrupts the Lipid metabolism regulation in placenta

We analyzed the gene expression of Lipid metabolism regulation genes. Gene expression of *Mtor, P53, Stat1, Stat3, CD36, DKK1, HMOX1* and *SRBI* of IVF group were significantly (p < 0.05) down-regulated, *TNF* and *PGF* were significantly (p < 0.05) up-regulated at E18.5 (Figure 8A). *Mtor, CD36, DKK1, PGF, Pth1h* and *SRBI* were significantly (p < 0.05) down-regulated, *P53, Stat1, Stat3, TNF* and *DGKK* were significantly (p < 0.05) up-regulated at E14.5 (Figure 8B).

**Figure 8.**
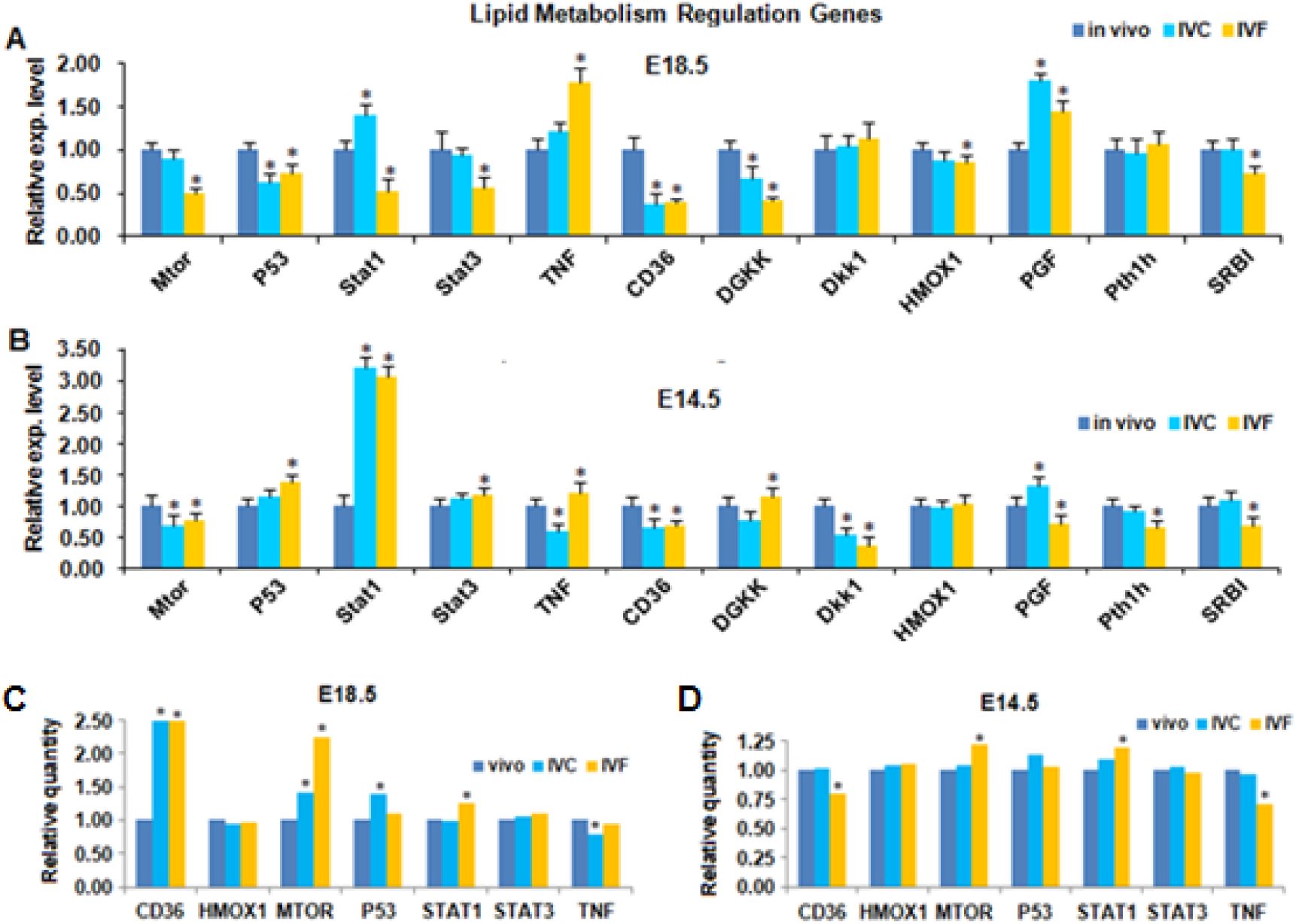
Expression of lipid metabolism regulation genes in the placenta. **A.** The gene expression levels of the three groups at E18.5. **B.** The gene expression levels at E14.5. **C.** Comparison of the protein quantity of lipid metabolism regulators (CD36, HMOX1, MTOR, P53, STAT1, STAT3, TNF) at E18.5 placenta of the three groups. **D.** The protein quantity results at E14.5 placenta. Values expressed as means ± standard errors. * p < 0.05 compared with control.

At protein level, expression of CD36, Mtor and STAT1 of IVF group were significantly (p < 0.05) up-regulated at E18.5 (Figure 8C). Expression of CD36 and TNF of IVF group were significantly (p < 0.05) down-regulated; MTOR and STAT1 were significantly (p < 0.05) up-regulated at E14.5 (Figure 8D). So IVF changed the expression of these genes in Mtor pathway which is the key pathway for lipid metabolism.

### 5 IVF leads to aberrant methylation and expression of imprinted genes related to lipid metabolism

Many imprinted genes participated in placenta lipid metabolism. We screen 32 imprinted genes from two imprinted genes databases: Mousebook and IGC-UOtago. Firstly, we test the methylation status of two key imprinted genes as *GANS, Grb10*. *GNAS* is a paternal imprinted genes and expressed in placenta. At E18.5, total methylation level of *GNAS* ICR region of IVF group is significantly higher than *in vivo* group (Figure 9A), but lower than in vivo group at E14.5 (Figure 9B). For the detail methylation level of the CpG sites on *Gnas* ICR region, please see Figure S1. *Grb10* is also paternal imprinted. The methylaton status of *Grb10* in IVF group is similar to *GNAS.* Compared to *in vivo* group, methylaton status of *Grb10* in IVF group is significant higher at E18.5 (Figure 9C), and is lower at E14.5 (Figure 9D). For the detail methylation level of the CpG sites on *Grb10* ICR region, please see Figure S1.

**Figure 9.**
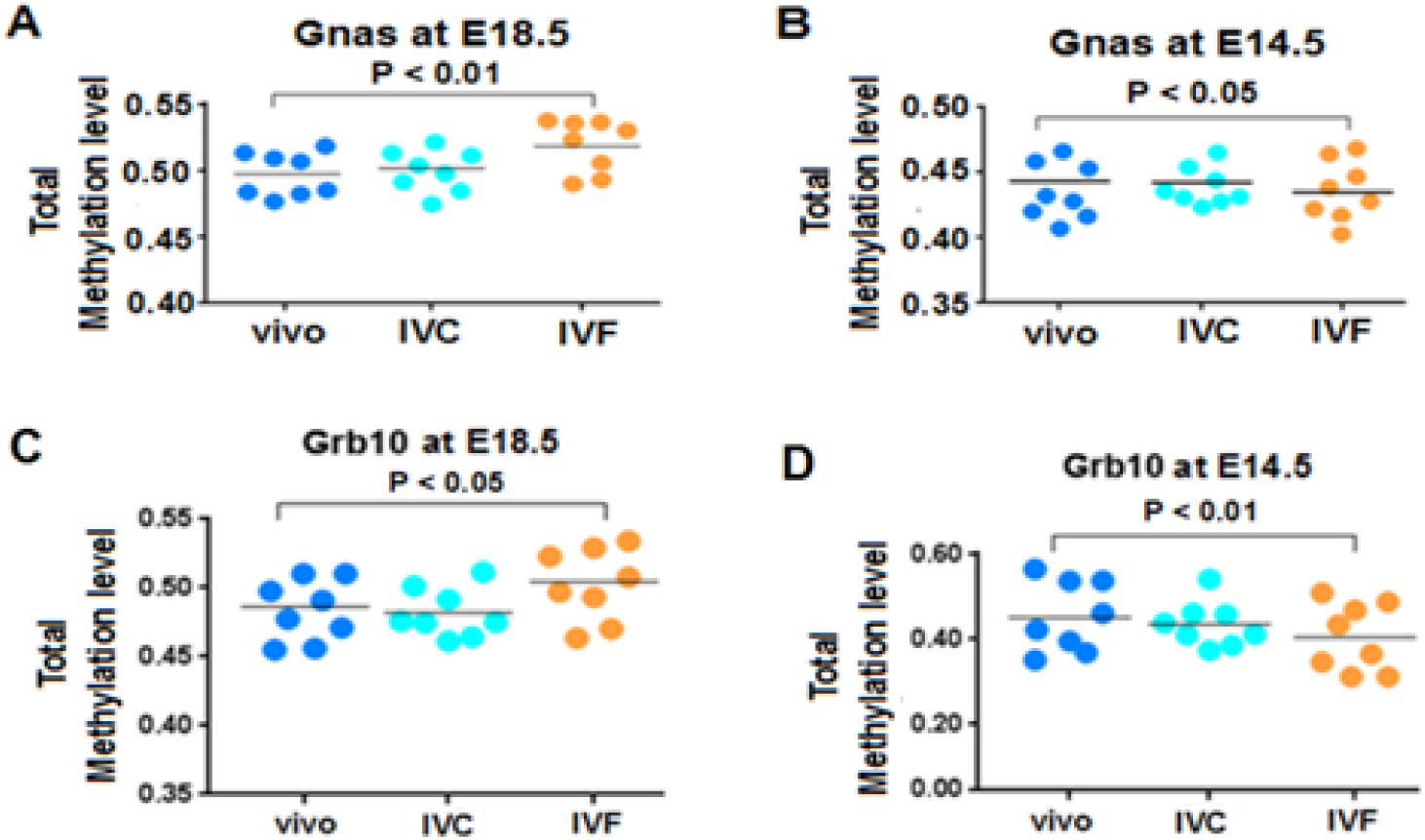
DNA methylation level of *Gnas* and Grb10 in the placenta. **A.** Total methylation levels of *Gnas* ICR region at E18.5. **B.** Total methylation levels of *Gnas* ICR region at E14.5. **C.** Total methylation levels of *Grb10* ICR region at E18.5. **D.** Total methylation levels of *Grb10* ICR region at E14.5.

To the 32 imprinted genes, 15 ones belong to paternal imprinted genes and 17 ones belong to maternal imprinted genes. At E18.5, maternal imprinted genes *Airn, Dlk1, GAB1, IGF2, NAP1L5, PEG1, PEG10, Rtl1, Stmbt2, USP29* and *Zdbf2* are down-regulated compared to *in vivo* group; *DDC, PEG3* and *SNRPN* are up-regulated (Figure 10A). Paternal imprinted genes *COB1, COPG2, GATM, GRB10, H19, Klf14, MEG3, Slc22a3, Slc22a18* and *Zim1* are down-regulated compared to *in vivo* group; *Slc22a2* is up-regulated (Figure 10B).

**Figure 10.**
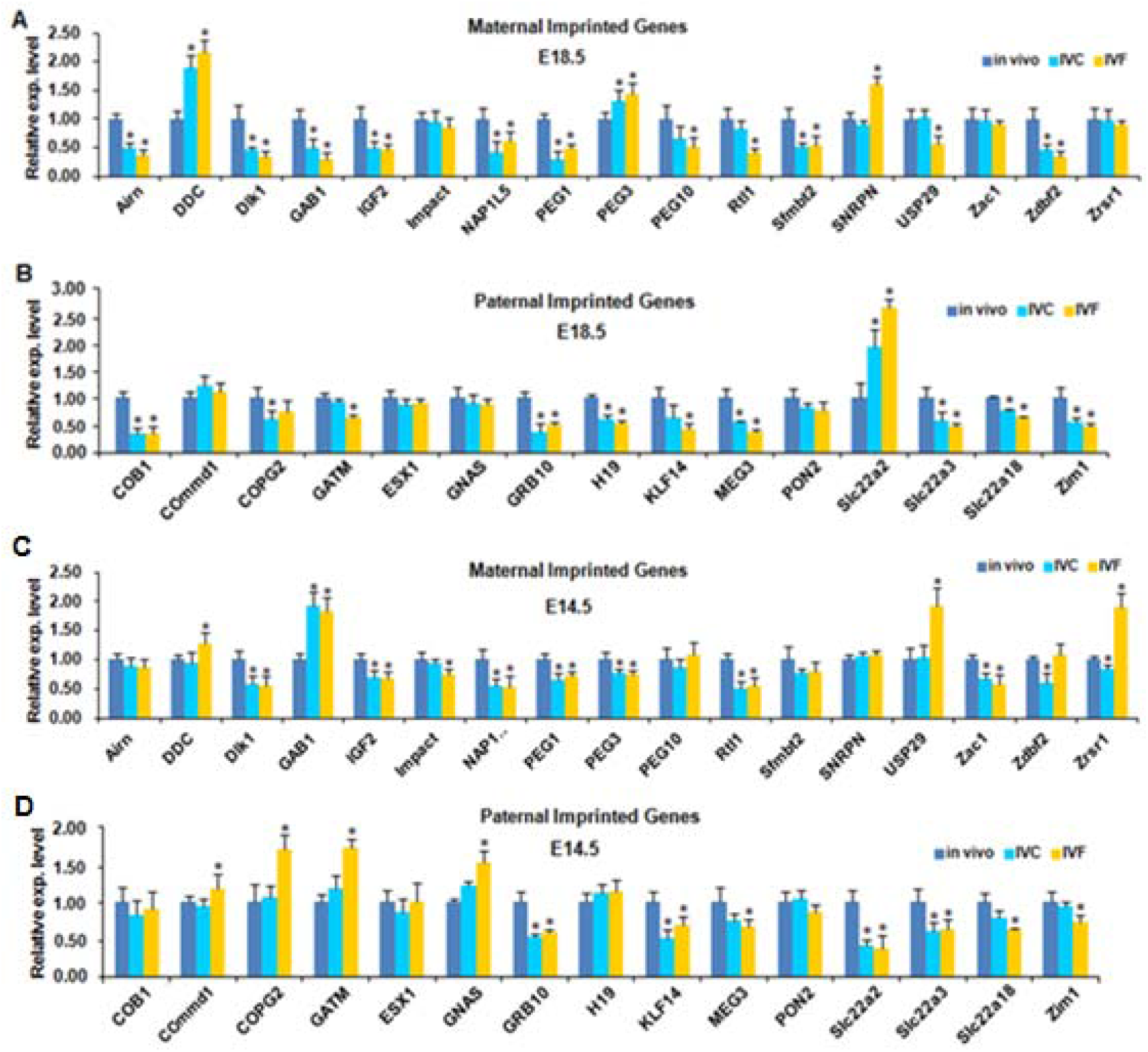
IVF resulted in aberrant expression of imprinted genes for lipid metabolism in placenta. **A.** Expression level of the maternal imprinted genes (*Airn, DDC, Dlk1, GAB1, IGF2, Impact, NAP1, NAP1L5, PEG1, PEG3, PEG10, Rtl1, Sfmbt1, SNRPN, USP29, Zac1, Zdbf2, Zrsr1*) at E18.5. **B.** Expression level of the paternal imprinted genes (*COB1, Commd1, COPG2, GATM, ESX1, GNAS, GRB10, H19, Klf14, MEG3, PON2, Slc22a2, Slc22a3, Slc22a18, Zim1*) at E18.5. **C.** Expression level of the maternal imprinted genes (*Airn, DDC, Dlk1, GAB1, IGF2, Impact, NAP1, NAP1L5, PEG1, PEG3, PEG10, Rtl1, Sfmbt1, SNRPN, USP29, Zac1, Zdbf2, Zrsr1*) at E14.5. **D.** Expression level of the paternal imprinted genes (*COB1, Commd1, COPG2, GATM, ESX1, GNAS, GRB10, H19, Klf14, MEG3, PON2, Slc22a2, Slc22a3, Slc22a18, Zim1*) at E14.5. Values expressed as means ± standard errors. * p < 0.05 compared with control.

At E14.5, maternal imprinted genes *Dlk1, IGF2, Impact, NAP1L5, PEG1, PEG3, Rtl1* and *Zac1* are down-regulated compared to *in vivo* group; *DDC, GAB1, USP29* and *Zrsr1* are up-regulate (Figure 10C). Paternal imprinted genes *GRB10, Klf14, MEG3, Slc22a2, Slc22a3, Slc22a18* and *Zim1* are down-regulated compared to *in vivo* group; *Commd1, COPG2, GATM, GNAS* are up-regulate (Figure 10D).

### 6 IVF resulted in aberrant gene expression and protein level of *DNMTs and Tets* in placenta.

*DNMT1, DNMT3A* and *DNMT3L*is important to maintain the methylation status of imprinted genes. Here we test the gene expression of these genes. Compared to *in vivo* group, *DNMT1, DNMT3A, DNMT3L, Tet1, Tet2* and *Tet3* of IVF group is down-regulated; *DNMT3B* is up-regulate (Figure 11A). At 14.5 placentae, *DNMT1, DNMT3A, DNMT3B, DNMT3L, TET1, TET2* and *TET3* are down-regulated (Figure 11B).

**Figure 11.**
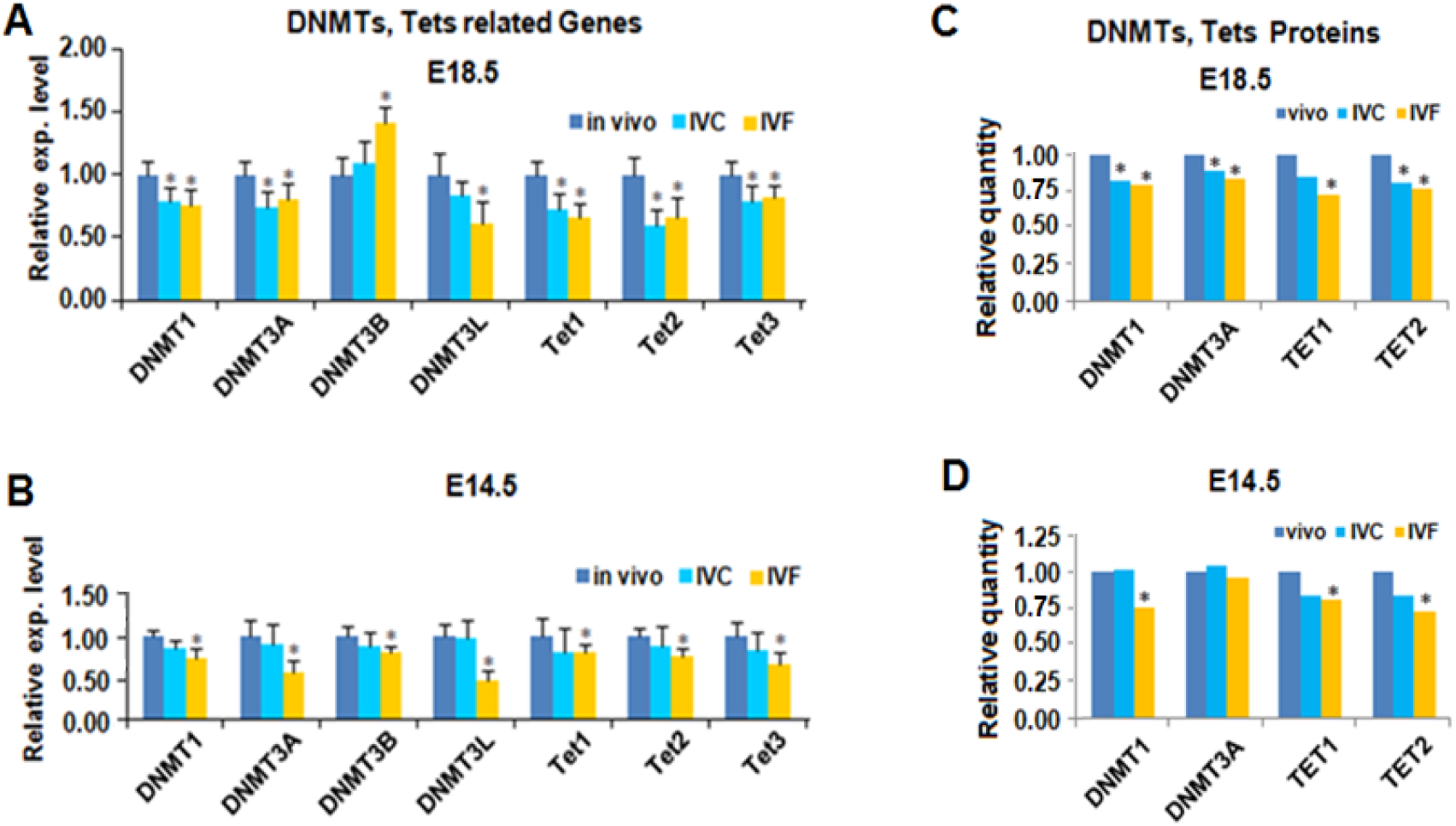
IVF resulted in aberrant gene expression and protein level of DNMTs and Tets in placenta. **A.** Gene expression levels of *DNMTs, Tets* at E18.5. **B.** Gene expression levels of *DNMTs, Tets* at E14.5. **C.** Protein quantity comparison of DNMTs, Tets at E18.55, respectively. **D**. Protein quantity comparison of DNMTs, Tets at E14.5. Values expressed as means ± standard errors. * p < 0.05 compared with control.

For protein relative quantification, DNMT1, DNMT3A, TET1, TET2 of IVF group are down-regulated compared to *in vivo* group at E18.5 (Figure 11C); at E14.5 DNMT1, TET1, TET2 are also down-regulated (Figure 11D).

### 7 Gene network for lipid metabolism in placenta

We sketched the cascade of lipid metabolism regulation network in mouse placenta Using Genemania [10]. *DNMTs* and *Tets* maintain the imprinted status of imprinted genes. Imprinted genes such as *GNAS, Grb10* participated in lipid metabolism with other regulators such as *Mtor, Stat1, Fasn* and *Cdc36*. *Lrp1, Apoc1, Ldlr, Plin2* and etc. are the genes for concrete lipid metabolism (Figure 12).

**Figure 12.**
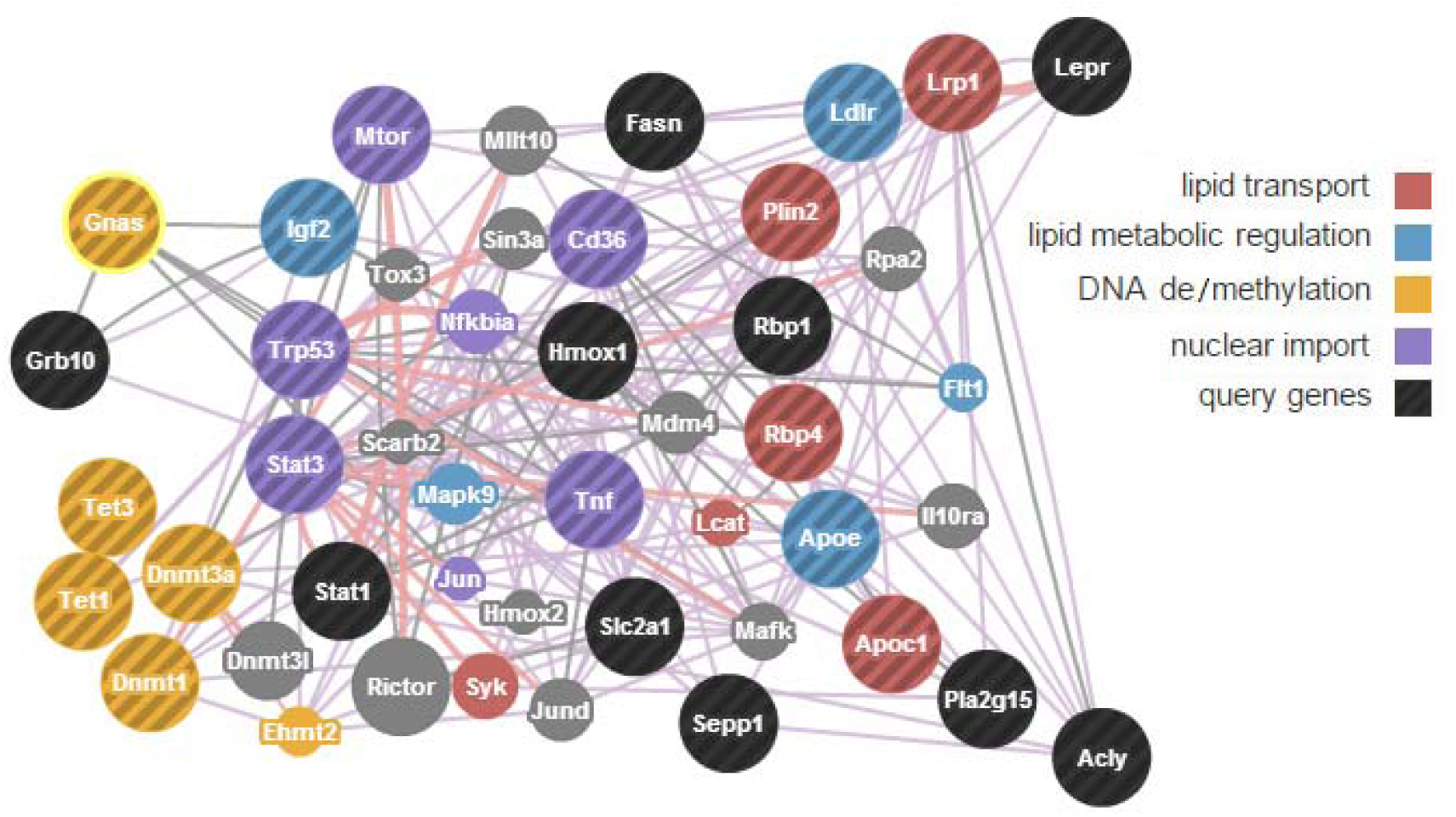
*GNAS* and the lipid metabolism regulation network in mouse placenta. There genes with important functions as lipid transport, DNA methylation and de-methylation, regulation of lipid metabolic are formed in a cascade pathway. Enzymes DNMTs and Tets modified the methylation status of *GNAS* and other imprinted genes. *Gnas* interact with *IGF2, P53, Mtor, Stats,* further regulate *Fasn, HMOX1, TNF, Slc2a1, Plin2, Lrp1, Apoe;* which participated in lipid metabolism.

## Discussion

The population of LBW will become large and large as more and more children born by ART worldwide. To birth weight, the body lipid or fat is take a big proportion as 50% dry mass for human [11], 40% dry mass for mouse [12]. The body lipid is also closely associated with the health status of individuals [13]. Human placenta and mouse placenta are in similar structure and belong to the same type: hemochorial placenta. So mouse is an optical animal for studying placenta.

Placenta efficiency is a micro-index for placenta function. The IVF group is significantly lower than *in vivo* group which indicate the nutrient transport capacity of IVF group is lower. “Bigger placenta, smaller fetus” is not only in mouse but validated by the epidemiological investigate [14, 15]. The IVC result is between that of IVF and in vivo. We excluded the litter number effect by transfer the same number blastocysts to every uterus.

LBW is accompanying by preterm birth or intrauterine growth retardation (IUGR). IUGR is more related to lipid metabolism than to Carbohydrate or proteins. As the lipid function of placenta is more complex which composed by uptake, transport and metabolism, the regulation of placenta Lipid function is also in different layers: imprinted genes, metabolism regulation genes, hormones and so on. The placental malfunction also related to preeclampsia, endometritis [16, 17]. Accumulated lipids in IVF placenta is the direct cause of LBW for IVF group.

The lipid metabolism is including β-oxidation, synthesis, combination, decomposition etc [18]. For cholesterols and acylglycerols is via facilitated transport from placenta to fetus. To another two type lipid phospholipids and sphingolipids, will be via active transport by transporters accompany by decomposition and synthesis. The lipidomic result shows the quantity of 15 lipid classes of IVF is 1.1-2.7 fold of in vivo group in E18.5 placenta. At E14.5, it is 0.2-1.0 fold of the in vivo group. So the placenta lipid metabolism is seriously affected by IVF. Placenta lipid uptake was affected by uterus environment like hormone, the nutrient of mother. If mother have diabetes and preeclampsia, the lipid support to fetal will be tighten [5, 17]. *Mtor* pathway is the major regulators for lipid transport [19]. If the transport capacity from placenta to fetal is limited, there will be a lot lipid accumulation as show in this study. Lipid accumulation will also change the morphology of placenta [20]. Other regulators are *Apoe, LRPs, PLA2Gs* and so on [21]. *Mtor, LRP8 and PLA2G10* is down-regulated in this study. IVF restrict the lipid transport capacity of placenta. *Mtor* pathway also regulated the lipid metabolism with *PTHrp, PPARs, STATs* and *IGF2* [21]. *ABCG1, CTP1A, EL, FASN, LIPIN1, LIPIN3* and *LPL are* down-regulated at E18.5; *APOC1, CTP1B* and *PPARA* are up-regulated at E18.5. IVF cause these gene expression spectrum indicate IVF cause lipid metabolism abnormal.

Imprinted genes are essential for placenta development and function. So the placental Mammals have evolution imprinted genes and balanced the nutrient form mother to fetus [22]. Paternal imprinted genes regulated more nutrients from mother to fetus, and maternal imprinted genes are prone to conserved nutrient in mother body [23]. This balance maintains a healthy development for fetal in uterus until to birth and the mechanism also foster new species by reproduction separation [24, 25].

*Gnas i*s the main regulator foe lipid metabolism in placenta [26-28]. *Gnas* knockout mice is fetal lethal with a defect placenta [29]. *Grb10* is paternal imprinted and function as a insulin receptor binding protein. It is specific expression in placenta and brain [30]. *Grb10* defect is related small placenta, low placenta efficiency, and LBW [31]. The biochemistry function of *Grb10* is promoting lipid decomposition and transport [32, 33]. To the paternal imprinted genes, such as *Dlk1* could regulate lipid metabolism [34]. It is with lower methylation level in IUGR mouse placenta [35]. *Dlk1* Knockout mice are with LBW, smaller placenta labyrinth layer [36]. The normal function of imprinted genes needs a normal imprinted status (methylation level). Both hypomethylation and hypermethylation will deteriorate its function. ART will trigger abnormal methylation of imprinted genes is very common [37, 38], and caused a higher mis-carrier rate than natural conceived pregnants [39, 40].

From the lipidomics to imprinted status, The IVF placenta exhibited significant perturbations at mid-to-late gestation. Imprinted genes play a vital role in determining placental phenotype and function. Our work demonstrates that structural abnormalities and dysfunction of the placenta may at least be partly caused by perturbation of lipid metabolism. These results support our hypothesis: imperfect IVF condition jeopardizes the imprinting status of imprinted genes for lipid metabolism regulation, which in turn causes abnormal expression of genes for lipid metabolism and its regulation, resulting in disruption of lipid metabolism, such as lipid accumulation and transportation, and thus contributes to low lipid transport efficiency leading to restricted fetal growth and LBW. Our study provides new insight to improve IVF procedures.

## Materials and Methods

### Animals

6-8 weeks ICR (CD-1) female mice, adult ICR male mice; Kunming vasectomized male mice for the pseudopregnant, Kunming adult female mice as surrogate mice. Mice were taken care in cages in the experiment animal center. It is under a constant 12h to12h light/dark cycle at 21–23°C room temperatures with free access to standard chow and tap water.

### Groups design

Female mice were superovulated and assigned to control group, IVF group and IVC group. Control group: blastocysts were collected after in vivo fertilization and then transfer to surrogate mice for in vivo development. IVC group: 1-cell embryos were harvested from the oviducts after in vivo fertilization and culture to blastocysts, then transfer to surrogate mice. IVF group: blastocysts were obtained after in vitro fertilization and culture, then transfer to surrogate mice. The three groups are all needed embryo transfer.

### Superovulation

Female mice were superovulated by intraperitoneal injection of 7.5 IU (0.15 ml) Pregnant Mare Serum Gonadotropin (PMSG, ProSpec, Israel), 48 h later by an intraperitoneal injection of 5 IU (0.1 ml) of human chorionic gonadotropin (HCG, ProSpec, Israel).

### In vitro fertilization and culture

13-15h later after the HCG injection, sperms were collected from the cauda epididymis of adult male ICR mice; Oocytes collected from oviduct ampulla of the female mice. In vitro fertilization was conducted in human tubal fluid (HTF) medium. Sperms were put in the incubator for capacitation. Capacitation of the spermatozoa was achieved by keeping them in a 37°C environment under 5% CO2 and 95% humidity for 1.5 h. The preincubated, capacitated sperm suspension was gently added to the dish of freshly ovulated cumulus-oocyte complexes, a final motile sperm concentration is opt to be 1–2 × 10^6^/ml. Sperm and oocytes were co-cultured for 8 h in insemination medium. In vivo or in vitro fertilized eggs (22–23 h after HCG), selected by the presence of two pronuclei, and then cultured under optimized culture conditions (KSOM+AA, Millipore). The developmental efficiencies in vitro were assessed by morphology under microscope.

### Embryo transfer recipients

Kunming females at least 6 weeks old were mated with vasectomized Kunming males 4 days prior to embryo transfer. The morning after mating, females were checked for the presence of vaginal plug, if it had then it was assumed to be on day 0.5 of pseudopregnant. Embryos were then transferred to the uteri of the pseudopregnant females on pseudopregnant day 3.5 according to standard procedures.

### Lipidomics analysis

3 placentae from three litters were pooled together in PBS then homogenization by Polytron (Turrax, IKA). Lipids were extracted using the optimized Bligh and Dyer’s method [41]. In detail, 750 ml of ice cold chloroform: methanol (1:2, v/v) was added to homogenized tissue suspended in 100ul of PBS. The mixture was vortexed vigorously for 1 min. After shaking at 1200rpm at 4 °C for 1 h, 250 ml ice cold chloroform and 350 ml of ice cold water was added to the samples. The samples were then subjected to another 1 min vortexing. The phases were separated by centrifugation at 9000 rpm for 2 min and the lower organic phase containing the lipids was transferred to a fresh tube. Lipids were re-extracted from the remaining aqueous phase with 500 ml of ice cold chloroform. The two extracts were pooled and vacuum-dried. Dried lipid film was then reconstituted in chloroform: methanol 1:1 (v/v) prior to analysis by mass spectrometry. The lipid samples were spiked with a mixture of internal standards prior to mass spectrometry. The standard cocktail contained -PA 17:0/17:0, PE 17/17:0, PG 14:0/14:0, d31-PI 18:1/16:0, PS 14:0/14:0, PC 14:0/14:0, C17 ceramide, C8 glucosylceramide, d6-cholesterol, d6-cholesterol ester and SM 18:1/12:0 [42]. Quantification of individual polar lipid species was performed using an Agilent 1200 HPLC system coupled with an Applied Biosystem Triple Quadrupole/Ion Trap mass spectrometer 3200Qtrap (Applied Biosystems, Foster City, CA) as described previously [43]. Mass spectrometry data was recorded separately under both positive and negative modes.

### RNA extraction and real-time PCR analysis

Pooled 6 placentae which obtained from at least two litters from the same group were used for RNA extraction. RNA was extracted with Trizol (Invitrogen Life Technology) according to the manufacturer’s instructions and treated with DNase I to eliminate genomic DNA contamination. Reverse transcript was accomplished using first-strand cDNA synthesis kit (QIAGEN). The RT reactions were performed to reverse transcribe 1 µg of total RNA. Real-time PCR was performed with the Bio-Rad CFX96 real-time PCR instrument (Bio-Rad, Hercules, CA) and SYBR Green ReadyMix for Q-PCR according to the manufacturer’s protocol (Sigma-Aldrich). Housekeeping gene GADPH is the reference gene for gene expression normalization. The primer sequences for the genes analyzed are listed in Table S2, Table S3 and Table S4, Table S6. Primer sequences were obtained from Primer Bank or designed using Primer Express. Expression level were analyzed using the ^△△^Ct method.

### Protein isolation and mass spectrum quantification

Three Placentae of the same group was pooled and homogenization by Polytron (TURRAX, IKA). Total Protein was isolated by CWBIO Kit (CW0891, Beijing). Protein concentration was assayed by BCA methods (Pierce Kit). Then the samples were treated by TMT Kit (Thermo Scientific) for Tandem mass tags. After the TMT tags were added, the samples were analyzed by HPLC (AKTA, GE). There are total 45 vials. Then three vials merged to one vial as No.1, 16, 31, then 2, 17, 32 and so on. The samples were dried in vacuum freeze drier (Labconco). When samples were dried, add 0.1% formic acid in it, storage at 4℃. 15 vials were put in the sample holder of Mass Spectrum (Thermo Finnigan LTQ). Then every sample was analyzed on Mass Spectrum. The raw data was used Mascot database for sequence alignment. Results were export as excels and used for data mining.

### DNA extraction and MassARRAY

Genomic DNA was extracted from placentae with Tissue DNA Kit (QIAGEN, Dusseldorf, Germany) according to manufacturer’s instructions. Absorbance at 260 and 280 nm was tested for DNA concentration and purity. The Sequenom MassARRAY platform (BGI, Shenzhen, China) was used to perform the quantitative methylation analysis. PCR primers were designed using the EpiDesigner software (Sequenom). The primer sequences are shown in Table S5. Epityper software v1.0 was used for calculating methylation ratios (Sequenom, San Diego, CA).

### Statistic analysis

Quantitative data is expressed as Means ± Standard deviation (SD). The data were subjected to ANOVA test for assessing any significant difference among the three groups. The least significant difference (LSD) post hoc test was used to examine any significant difference between groups. Lipidomic data were normally distributed and the Analysis of statistical significance of differences in molar fractions between the three groups was sought using Student’s two-tailed unpaired t-test. Probabilities lower than 0.05 were considered significant. Analyzes were performed using GraphPad Prism 5 software.

### Animal Ethics

This study was performed in strict accordance to the Guide for the Care and Use of Laboratory Animals of the NIH. All animal protocols were approved by the Committee on the Ethics of Animals and Medicine, the Institute of Genetics and Developmental Biology, Chinese Academy of Sciences.

## Supporting Information

**Figure S1** Methylation status of CpG sites of *Gnas* and *Grb10*.

**Table S1** The quantities of 15 lipid classes in mouse placenta.

**Table S2** qPCR primer sequences for the genes related to lipid uptake, transport and metabolism.

**Table S3** qPCR primer sequences for the genes related to lipid metabolism regulation.

**Table S4** qPCR primer sequences for the imprinted genes.

**Table S5** Sequenom MassArray primer sequences for *Gnas* and *Grb10*.

**Table S6** qPCR primer sequences for *DNMTs* and *Tets*.

## Author Contributions

Conceived and designed the experiments: YBW, FZS. Performed the experiments: YBW, SQC, SML. Analyzed the data: YBW, SML, FZS. Contributed reagents/materials/analysis tools: YBW, XYH, GHS. Wrote the paper: YBW, FZS.

